# *Toxoplasma gondii* establishes an extensive filamentous network consisting of stable F-actin during replication

**DOI:** 10.1101/066522

**Authors:** Javier Periz, Jamie Whitelaw, Clare Harding, Leandro Lemgruber, Simon Gras, Madita Reimer, Robert Insall, Markus Meissner

## Abstract

Apicomplexan actin is well conserved and clearly important during the parasite's life cycle. Several studies assert that its polymerization kinetics are unusual, permitting only short, unstable F-actin filaments. However, it has not been possible to study actin *in vivo*, so its physiological role has remained obscure. This has led to functional models which are mutually conflicting, incompatible with actin behavior from other eukaryotes, and cannot explain actin's importance during basic processes such as parasite replication and egress. Here we use a chromobody that specifically binds F-actin to demonstrate that *Toxoplasma* forms stable actin filaments *in vivo*. F-actin is not only important for parasite replication, but also forms an extensive network that connects individuals both within and between parasitophorous vacuoles, and allows vesicles to be exchanged between parasites within a vacuole. During host cell egress, prior to motility, this network collapses in a calcium-dependent manner. This study demonstrates unexpected roles of *Toxoplasma* actin during the asexual life cycle, and proves that formation of F-actin depends on a critical concentration of G-actin, implying a polymerization mechanism similar to mammalian actin.

**One Sentence Summary:** *Toxoplasma* establishes a stable F-actin network that is essential for replication and material transport between individual parasites.

## Main Text

For the past 20 years, biochemical assays using heterologously expressed apicomplexan actin have implied that it only forms very short filments that are intrinsically unstable (*1–3*). Furthermore, it is believed that it polymerizes in a highly unusual, isodesmic manner (*4*) and it has been suggested that this represents an important adaptation to provide the force for apicomplexan motility. While recent studies also implicate parasite actin in other essential functions, such as maintenance of the apicoplast, a plastid-like organelle (*5, 6*), daughter cell replication (*7*) or motility of secretory organelles (*8*), these functions would predict the polymerization of longer filaments. However, to date, it has not been possible to study actin *in vivo* and in real-time in order to correlate its assumed functions with the localization and dynamics of F-actin.

Here we expressed different F-actin binding proteins fused to fluorescent reporters and found that expression of actin chromobodies (*9, 10*), which consist of camelid nanobodies (*11*) fused C-terminally to a Halo tag (Cb-Halo) (*12*), (*13*), is tolerated well by the parasite and can be stably expressed without causing phenotypic abnormalities (Fig.S1). No differences could be observed in replication, egress or host cell invasion for parasites expressing Cb-Halo while gliding motility rates were slightly enhanced (Fig.S1), suggesting that expression of actin Cb-Halo has no significant effects on actin dynamics in the parasite.

We next localized Cb-Halo using a cell permeable halo ligand coupled to TMR both in live and fixed parasites. Surprisingly, individual daughter parasites remain connected at the posterior pole via actin filaments throughout intracellular development (Fig.1, Movie S11). With the increasing size of the parasitophorous vacuole (PV) these connections expand, resulting in the formation of an extensive network within the PV, which appears to originate from parasites localized closer to the center and extends to the surface of the PV (Fig.1A,B,3C, Movie S1). Incubation of wild-type parasites with antibodies preferentially binding to F-actin (14) revealed a similar organisation, though the network was only partially stained with this antibody (Fig. 1A, Fig.S2, Movie S2). We verified co-localization of Cb-Halo with alpha-actin in jasplakinolide (Jas) stabilized filaments (Fig.S2). Furthermore, incubation of parasites with the actin filament disrupting compound cytochalasin D (CD) resulted in a complete collapse of this network (Fig. 1A, Fig.S3 Movie S3), whereas stabilization of F-actin using Jas caused the formation of a thicker, more elaborate filamentous network (Fig.1A, Fig.S2, Movie S4). Upon addition of Jas, the cytosolic stain completely disappears and F-actin can be exclusively detected at the apical and polar ends of the parasite (Fig.S2). To investigate the specificity of Cb-Halo for parasite actin, we performed immunoblots under native conditions using lysates obtained from extracellular parasites. We find that Cb-Halo binds to approximately 5% of total actin (Fig.S2). Together these data confirm previous reports suggesting that 98% of actin is in its globular form in extracellular parasites (*1*).

**Fig. 1.**
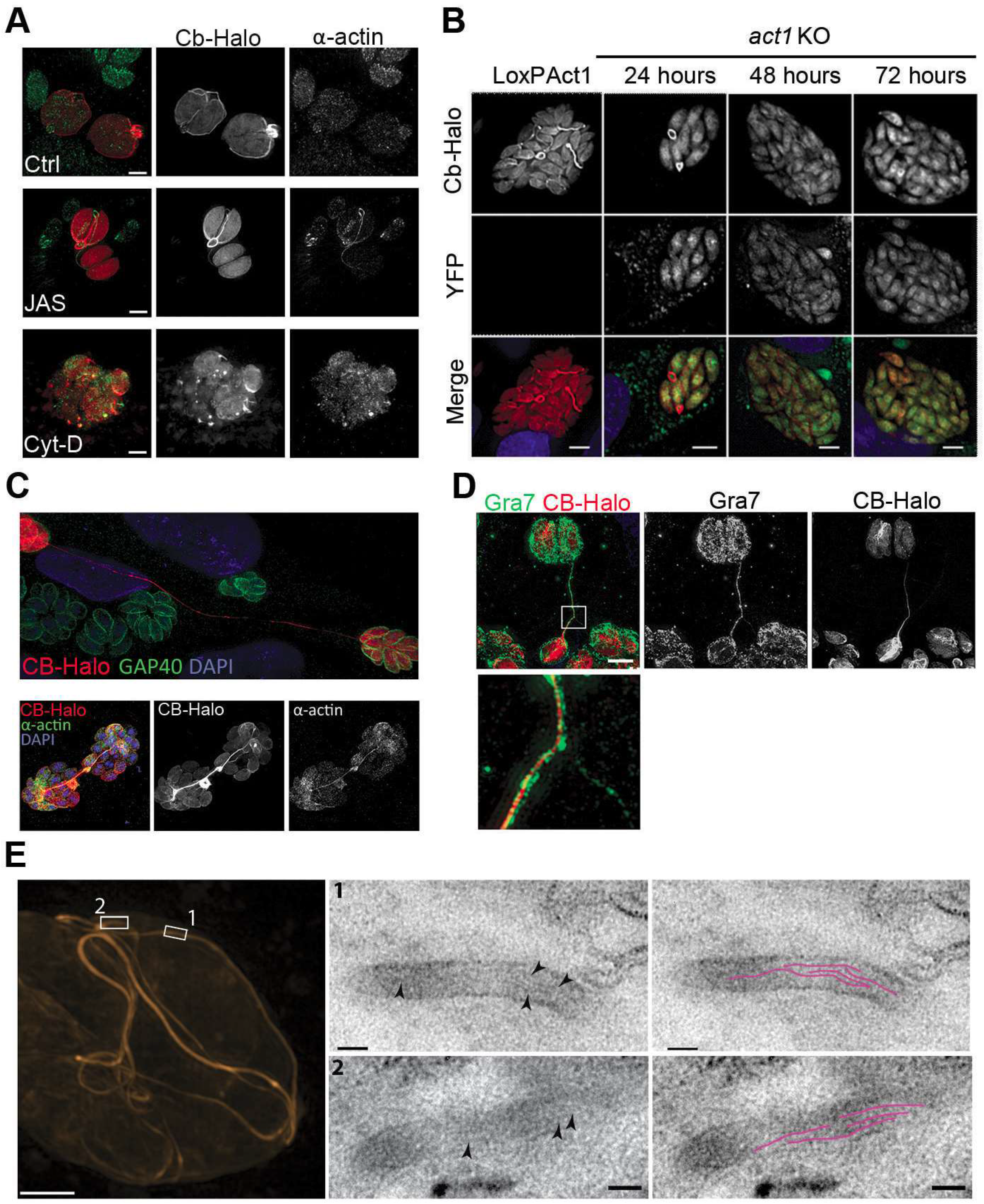
Filamentous actin can be visualized by expression of Cb-Halo. A) 3D imaging of parasites transfected with Cb-Halo and stained using a specific apicomplexan actin antibody (14). An actin filament network can be observed using Cb-Halo with only partial colocalization of Cb-Halo and actin antibody in filaments connecting parasites. Upon treatment with 2 μM CD, the filamentous network collapses and colocalization of Cb-Halo and actin antibody in regions of collapsed filaments is observed. Upon treatment with 100 nM Jas for 1 h, a more elaborate network can be seen with enhanced co-localization between the antibody and Cb-Halo. Scale bar 2 μm. B) Expression of Cb-Halo in a conditional *act1* cKO. A filamentous network is observed prior to excision of *act1* in loxPAct1. As soon as 24 hours after excision of *act1* the network diminishes. No filaments are observed at later time points. C) Long connections between vacuoles can be observed, some extending for over 50 μm. Scale bar 10 μm D) The dense granule protein Gra7 is seen surrounding Cb-HALO connections. However some Gra7-positive tubules are seen in the absence of F-actin. Scale bar 10 μm. E) A vacuolar network was imaged with SIM super-resolution microscopy and the same areas were imaged with TEM. Filaments of 5 nm in thickness were present within the network tubules, extending over 100 nm in length. Scale bars: 200 nm (3D-SIM); 50 nm (TEM).

Importantly, when we analyzed the localization of Cb-Halo in a conditional null mutant for *act1* (*act1* cKO) (*5*), we found that as early as 24 hours after removal of *act1*, the filaments collapsed, similar to the effect observed with CD treatment, suggesting that at this time point G-actin is below the critical concentration required for formation of F-actin, strongly arguing against an isodesmic polymerization process (*4*). In good agreement, at later time points, a filamentous network was not observed in *act1* cKO parasites (Fig.1B).

Surprisingly, we also detected very long (>50 μm) extensions that connect two independent vacuoles (Fig.1C). Examination of these long, extra-vacuolar filamentous connections shows that they were formed of an F-actin core surrounded by dense granule proteins, such as Gra2 and/or Gra7 (Fig.1D, Fig.S2). Intriguingly, we observed also connections devoid of F-actin, suggesting a dynamic behavior of F-actin during parasite development where it is possible that F-actin depolymerizes once connections are formed (Fig.1D, Fig.3C).

To investigate the presence of F-actin within this network, we performed correlative light-electron microscopy (CLEM). The network within a vacuole was imaged with super-resolution microscopy (3D-SIM) (Fig. 1E). Thin sections of the same network were imaged with transmission electron microscopy (squares). Within the network tubules, ~5 nm thick filaments (arrows) extending over 100 nm in length were observed (highlighted in magenta).

Together these data demonstrate that parasite actin forms stable F-actin filaments during intracellular development of the parasite and that – as is the case in mammalians and plants - actin polymerization depends on a critical concentration of available G-actin.

Next, we were interested in the stability and behavior of the F-actin network in live parasites. Using time-lapse analysis during parasite replication, we found that the filamentous network is dynamic and the length of filaments appears to depend on the cell cycle (Fig.2A, Movie S5). During replication of parasites, the network appears to collapse, adopting a ‘ring-like’ appearance (asterisks) before reforming and extending throughout the PV. However, when fluorescence recovery after photobleaching (FRAP) was used to estimate the turnaround time of G-actin within individual tubules, we were unable to detect any fluorescent recovery in established tubules, even after prolonged ob servation (longest time scale ~120s), suggesting that once F-actin structures are formed, they must depolymerize completely before new actin is incorporated (Fig.2B,C).

**Fig. 2.**
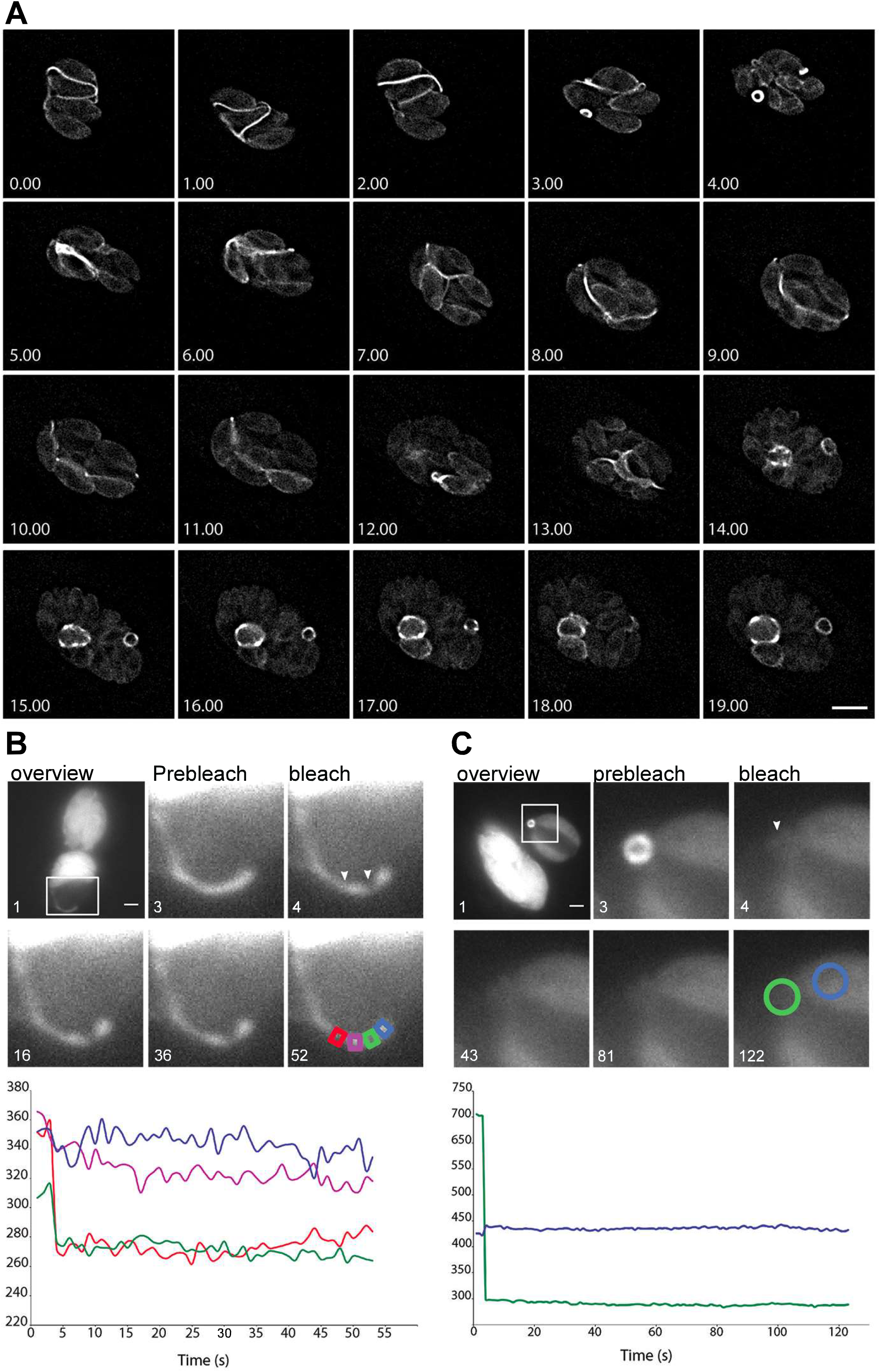
The F-actin network is dynamic during replication but individual filaments are stable. **A**) Analysis of Cb-Halo during two rounds of replication. Images were taken every 30 mins for 20 h. The network appears dynamic across the intracellular lifecycle, collapsing into rings during daughter cell emergence (asterisks). Scale bar 5 μm. Time lapse imaging of a filament (**B**) and the residual body (**C**) before and after photobleaching (white arrow) show no recovery of average fluorescence intensity, indicating that actin filaments were stable during imaging. Plot shows average fluorescence intensity (red and green squares in B and green circle in C) before and after photobleaching, compared with untreated areas (blue and magenta squares in B and blue circle in C). FRAP treatment shows a dramatic decrease in fluorescence intensity in the bleached area with no recovery of fluorescence during the duration of the experiment (52 and 122 s respectively), indicating the presence of stable actin filaments and a lack of significant turnover. Scale bar 2 μm.

These data suggest that actin turnover within the filamentous structures is slow in the absence of any cell-signaling that may trigger specific parasite events. We speculate that the observed dynamic behavior is exclusively caused by active extension and retraction of filaments at the posterior pole of the parasite. It remains to be seen if the tubular structures contain long actin filaments or, as might be assumed due to the reported properties of apicomplexan actin, short F-actin bundles that are connected via actin binding proteins.

Conditional deletion of *act1* is surprisingly well tolerated by *T. gondii* (*5*) and previous studies on parasite actin suggested only a minor role of actin during parasite division (*15*). However, closer examination of individual parasites within the PV revealed that while wild-type parasites form a conventional rosette, *act1* cKO parasites are not well organised, with erratic appearance. Actin filaments connect parasites at their posterior pole whereas no obvious connections between individual parasites were observed in the *act1* cKO (Fig.1B, 3A). Furthermore, a residual body could not be observed upon depletion of *act1* and parasites appeared to have aberrant morphology at the posterior pole, which appears in general flattened (Fig.3B).

**Fig. 3.**
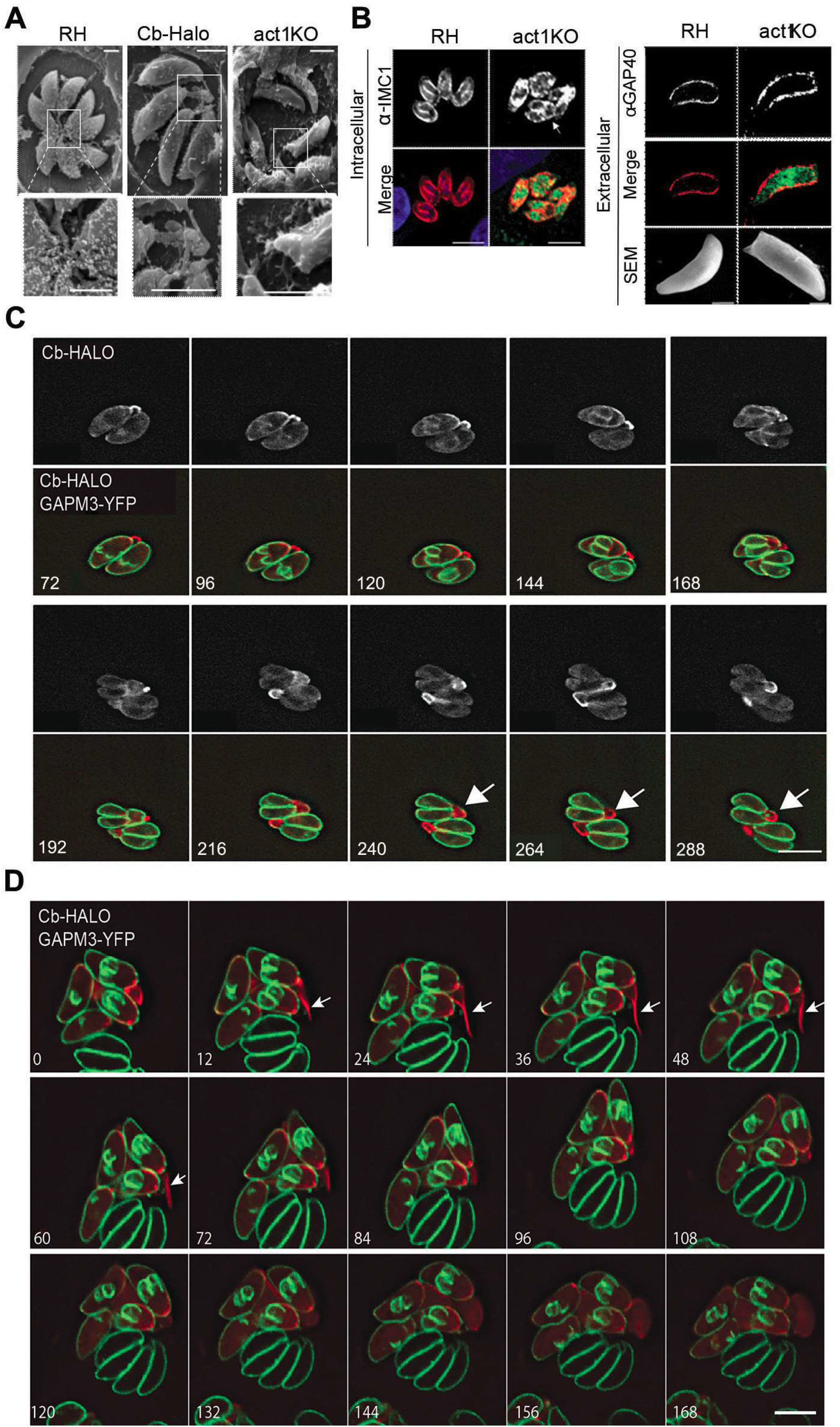
Analysis of F-actin dynamics during replication. **A**) Scanning EM of wild type, Cb-HALO and *act1* cKO parasites showing the filamentous network connecting parasites from the posterior end. *act1* cKO parasites are disorganized within the vacuole and form no connections. **B**) Analysis of the IMC of the *act1* cKO using IFA and scanning EM analysis in both intra- and extracellular parasites. Arrowhead shows the end of division of the *act1* KO. *act1* cKO parasites have a flattened bottom, torpedo shape. Scale bars: fluorescence images: 10 μm, SEM: 2 μm. **C**) GAPM3-YFP expressing parasites transiently expressing Cb-Halo were imaged every 6 mins for 5 hours, F-actin can be seen initially connecting the basal end of the parasites before accumulating beneath the forming daughter cells where it appears to concentrate towards the rear of the new daughters during emergence and recycling of the maternal IMC. Scale ar 5 μm. **D**) Another example of F-actin behaviour during replication using conditions as in (C). Note that in this example a F-actin connection is formed between two neigbouring PVs. The connection rapidly extrudes (arrow), remains in contact with the neighbouring PV for several minutes, before it retracts.

To analyze the role of F-actin during intracellular replication in detail, we performed localization analysis of Cb-Halo in parasites co-expressing endogenously tagged GAPM1a-YFP as marker protein for the inner membrane complex (*16*) (Fig.3C, Movie S5). During early stages of daughter cell formation, F-actin was found primarily at the posterior end of the parasite (Fig.3C), later F-actin accumulates at the nascent IMC of the daughter cells (Fig.3C, arrowhead) and further concentrates at the posterior end of the mother cell, where it colocalizes with the mother IMC (Fig.3C, arrow). Finally, towards the end of replication, the IMC of the mother collapses and individual GAPM1a-YFP positive vesicles are formed that localize with F-actin and are transported towards the daughter cells IMC. This suggests an important role of F-actin in recycling of mother cell IMC and the final assembly of the daughter cell IMC. Together with the phenotypic defects, this analysis demonstrates a function of parasite F-actin during formation of the IMC of daughter cells.

During this analysis we also identified events where F-actin connections are formed between PVs (Fig.3D). These connections are formed in a highly dynamic manner. Cb-Halo stained filaments were observed to rapidly extrude from one PV and remain in contact with a neighboring PV for a few minutes before retracting (Fig.3D, Movie S6).

Together these observations suggest an exciting hypothesis that individual parasites both within and between vacuoles may exchange material during intracellular development along these filamentous structures.

To test this hypothesis in more detail, we analyzed if intra-parasite material is indeed transported along the identified filaments. Time-lapse analysis of both the integral membrane protein GAPM1a-YFP and cytosolic GFP was performed in wt parasites (Fig.S3) and in both cases vesicles could be identified that moved outside parasites (Fig.S3, Movie S7). Digital tracking of vesicles correlated well with actin filaments described above (Fig.S3). By expressing Cb-Halo in these lines, we were able to show that these vesicles were closely associated with F-actin both in fixed (Fig.4A) and live cells (Fig. 4C,D). Importantly, movement of extracellular vesicles was F-actin dependent; incubation of parasites with CD significantly abrogated extracellular vesicular motility (Fig.4B). No motility could be observed in *act1* cKO parasites.

**Fig. 4.**
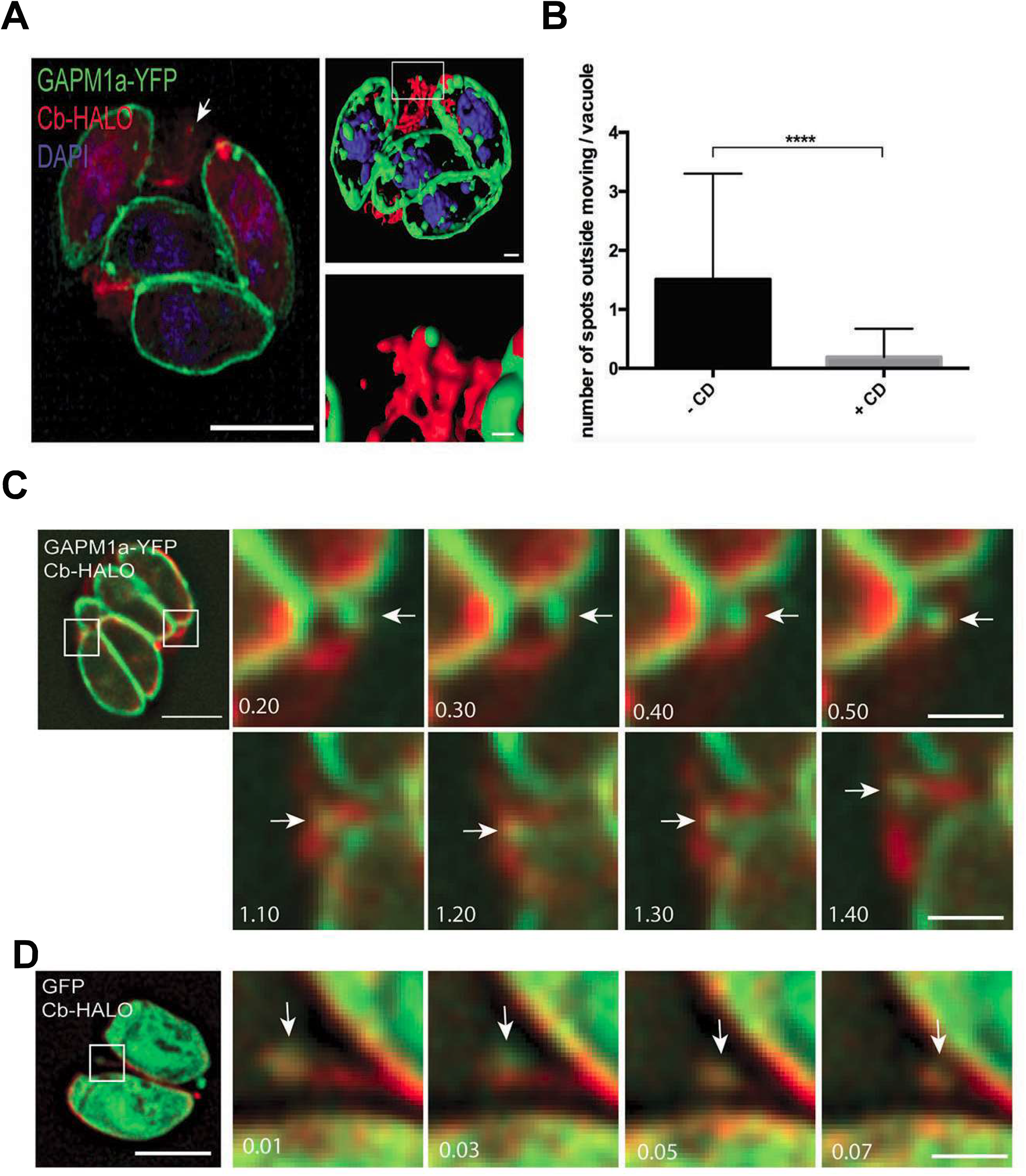
Parasite-derived extracellular vesicles are transported in an F-actin dependent manner. **A**) In parasites endogenously expressing GAPM1a-YFP, extracellular vesicles could be observed in close apposition to Cb-HALO labeled filaments (arrow). **B**) The number of vesicles per vacuole that moved within 5 min of imaging were quantified in the presence and absence of 500 nM CD. At least 60 vacuoles were counted over three independent experiments. **** P < 0.0001. Parasites transiently expressing Cb-HALO were imaged every second for up to 5 mins. Extracellular vesicles positive for GAPM1a-YFP (**C**) and GFP (**D**) were observed to move along F-actin filaments (arrows). Scale bar, 1 μm.

Finally we tested how the organization of the F-actin network changes during parasite egress, especially as *act1* cKO parasites are completely blocked in egress, while gliding motility and host cell invasion can still be observed in the absence of detectable F-actin (*5*). When we triggered a calcium signaling cascade using Ca^2+^-Ionophore (*17*), the F-actin network immediately collapsed. This collapse is rapid and can occur as quickly as 10 seconds after the addition of Ca^2+^-Ionophore even if the parasites do not egress from the cell (Fig.5A, Movie S8). Collapse of the network is followed quickly by the initiation of motility and egress from the host cell (Fig.5B, S. Movie S9). As the parasites begin to move, F-actin can be detected at the rear of the parasites. Importantly, this indicates that collapse of the network is required for egress and suggests a role in signaling prior to escape.

**Fig. 5.**
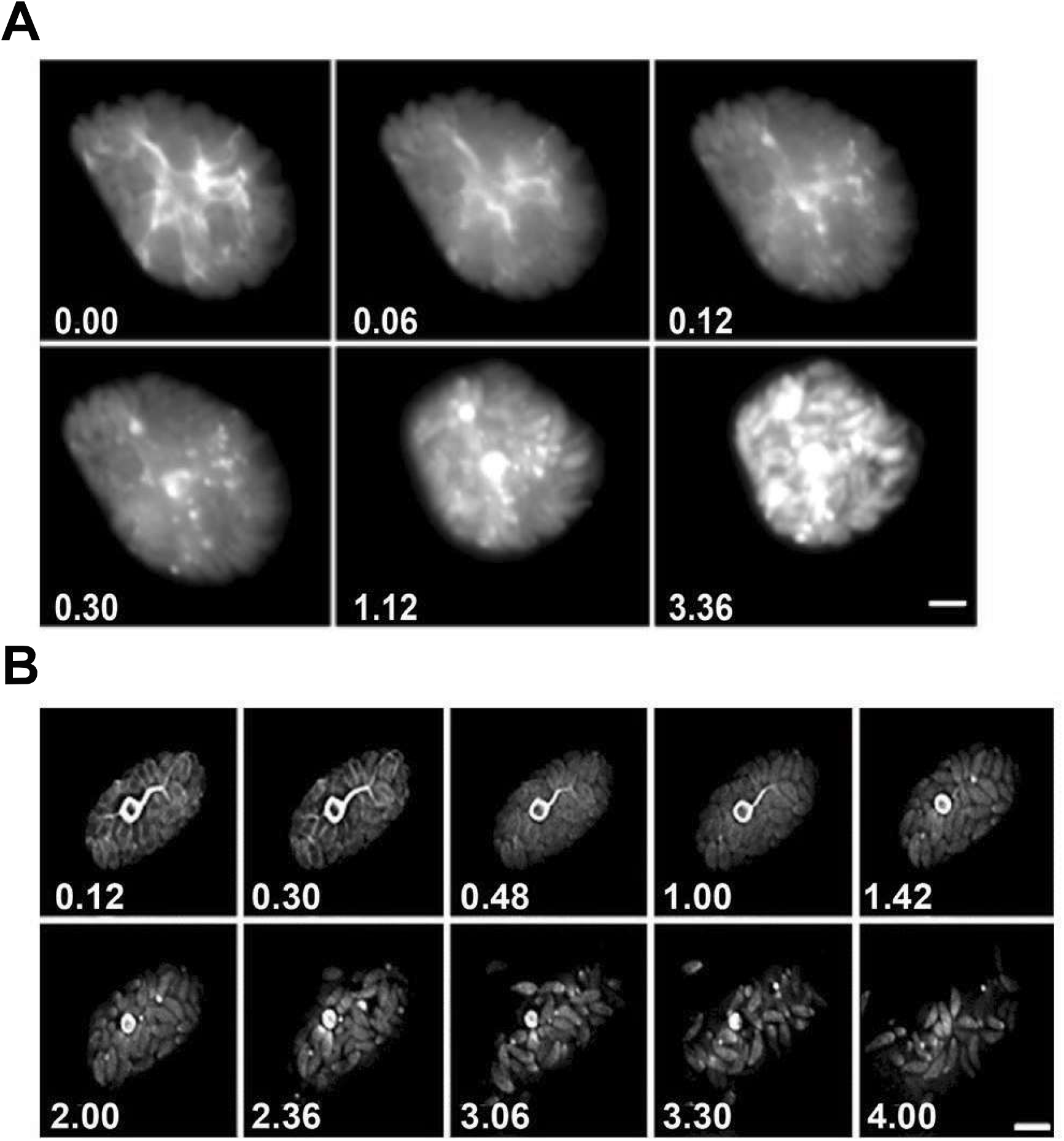
Collapse of the F-actin network can be triggered by calcium ionophore. **A**) A flattened Z-stack of the dynamics of F-actin after addition of Ca^2+^, images taken every second. Filaments break up in a calcium dependant manner even if the parasites do not egress. **B**) The network collapses before parasites begin to egress. While filaments quickly collapse, the residual body remains intact during egress. Scale bar, 10 μm.

In summary, we detect for the first time actin filamentous structures in apicomplexan parasites and demonstrate that *Toxoplasma* actin has essential and previously unforeseen roles during the intracellular development of the parasite (Fig. 6). Actin serves several functions: from segregation of the apicoplast; recycling of organellar material from the mother to the daughter cells, and is potentially required for structural stability of the vacuole and intracellular communication between individual parasites within the same or independent PV. In order to function, we show here that parasite actin is capable of forming stable actin filaments and that, (at least) *in vivo*, this polymerization depends on a critical concentration of G-actin. This first demonstration of the presence and functions of stable, filamentous actin in *Toxoplasma* offers new insights into the biology of apicomplexan parasites.

**Fig. 6.**
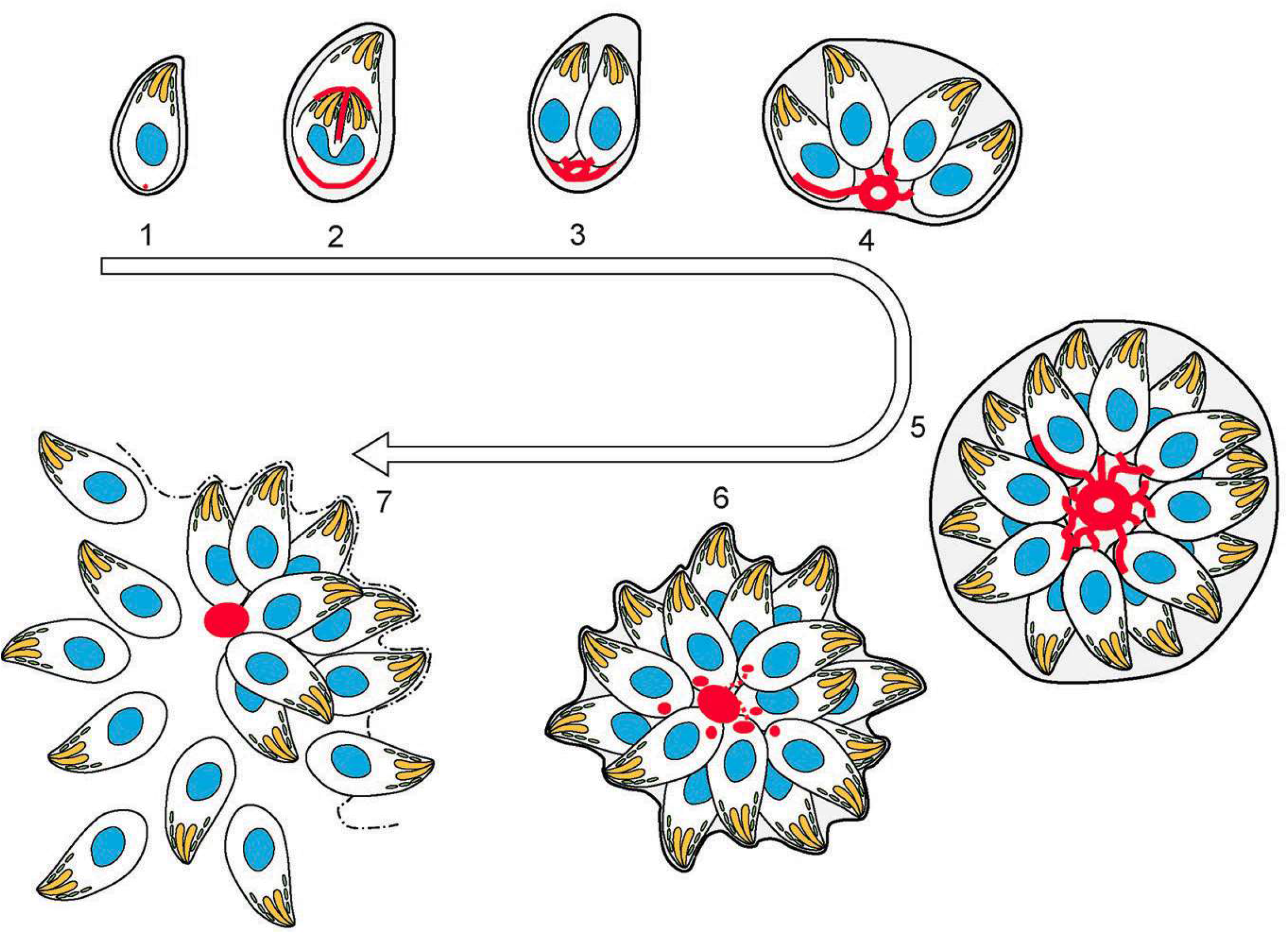
Representation of actin dynamics during replication. 1) After successful invasion, tachyzoites establishe a parasitophorous vacuole and start replication. 2) During daughter cells formation, actin labelling is observed at the developing IMC of the daughter cells and at the basal pole of the mother cell. 3) Once the daughter cells are fully formed, the actin signal strongly colocalises with the mother IMC, as it is recycled. 4-5) Replication continues an an extensive filamentous network is established between individual tachyzoites. 6) After treatment with a Ca^2+^ ionophore, filaments between parasites are desolved. The network collapses and dots of actin are detected at the posterior end of tachyzoites. 7) Tachyzoites egress from the vacuole leaving behing an actin accumulation in the residual body.

## Acknowledgments

We would like to thank Gary Ward (University of Vermont) for the IMC1 antibody, Jake Baum (Imperial College, London) and Dominique Soldati (University of Geneva) for Act1, GAP40 antibodies. Freddy Frischknecht and Isabelle Tardieux for critically reading the manuscript. Special thanks to Dr. Gurman Pall and other lab members for stimulating discussions. CRH is supported through a Sir Henry Wellcome Fellowship (WT103972AIA). This work was supported by an ERC-Starting grant (ERC-2012-StG 309255-EndoTox) and the Wellcome Trust 087582/Z/08/Z Senior Fellowship for MM. The Wellcome Trust Centre for Molecular Parasitology is supported by core funding from the Wellcome Trust (085349). The funders had no role in study design, data collection and analysis, decision to publish, or preparation of the manuscript.

The authors have declared that no competing interests exist.

## Supplementary Materials

Materials and Methods

Figures S1-S4

Movies S1-S9

